# Profiling of *NUMT*s in Gingivobuccal Oral Cancer

**DOI:** 10.1101/2020.11.13.381020

**Authors:** Arindam Palodhi, Tanisha Singla, Arindam Maitra

## Abstract

Very little information exists on the NUclear MiTochondrial sequences (*NUMT*s) in oral squamous cell carcinoma-gingivobuccal (OSCC-GB). We analysed whole genome sequences obtained from paired tumour and blood DNA from 25 patients of OSCC-GB for detecting *NUMT*s. Nine potential somatic *NUMT*s and 78 germline *NUMT*s were identified in these patients. None of the somatic *NUMT*s could be confirmed by PCR assay. None of the germline *NUMT*s were found to be specific for OSCC-GB. Although there have been recent reports on detection of somatic *NUMT*s in other cancer types, our results suggest *NUMT*s, both somatic and germline, are not involved in OSCC-GB.

## Introduction

The mtDNA fragments localized into the nuclear genome are identified as NUclear MiTochondrial sequences (*NUMT*s) and the processes which generate *NUMT*s are collectively known as *NUMT*ogenesis (Singh et al., 2017). Irrespective of the insertion site and the mtDNA sequence, all *NUMT*s found in human are rendered non-coding. This pseudogenization of *NUMT*s is due to the difference in codon usage between the nDNA and the mtDNA (Gellissen et al., 1987; Perna et al., 1996).

Although role of *NUMT*s in mitochondrial-nuclear genome evolution has been studied considerably, information on the *de-novo NUMT*s in both sporadic (Turner et al., 2003) and inherited diseases (Goldin et al., 2004) are scarce. Involvement of *NUMT*s in cancer remains a relatively underexplored yet important area of research.

## Study participants and methods

Twenty-five patients of gingivobuccal oral cancer (OSCC-GB) were recruited with voluntary informed consent (ICGC India Project Team, 2013). Blood samples collected with informed consent from seven non-cancer individuals were used as control samples.

Whole genome sequence data was generated from paired tumor and blood DNA of 25 OSCC-GB patients by paired end sequencing using Illumina Hiseq-2500. BWA-MEM (Li, 2013) was used to align the whole genome FASTQ files to the human reference sequence (hg19) + mitochondrial reference sequence (rCRS-revised Cambridge Reference Sequence). Alignment files thus produced were used as input for Discovery of Nuclear Mitochondrial Insertions (DINUMT; Dayama et al., 2014) to identify *NUMT*s. Circos plots shown in this study were generated using BioCircos (Cui et al., 2016).

For the PCR validation of the *NUMT*s, the individual primer pairs were designed in a manner, such that they would amplify the entire length of the *NUMT + at least 150 bases upstream + at least 150 bases downstream* of the insertion site and in absence of *NUMT*s they will amplify only the buffer region **(Supplementary Table S1, Supplementary Table S2)**.

## Results

*NUMT*s detected in both paired tumours-blood samples were defined as germline *NUMT (G-NUMT)* and *NUMT*s present only in tumours samples but not in paired blood DNA were defined as somatic *NUMT* (S-*NUMT*s). Using *DINUMT,* we could detect 50 such S-*NUMT*s in 25 OSCC-GB patients. Each of these S-*NUMT*s were visually inspected using IGV (Robinson et al, 2011). After verification in IGV, we found 9 potential S-*NUMT*s in 6 patients were supported by at least two discordant reads and one split read **(Figure 1A, Table 1)**. Only these 9 S-*NUMT*s which were supported by both discordant and split reads were investigated further.

**Figure 1:**
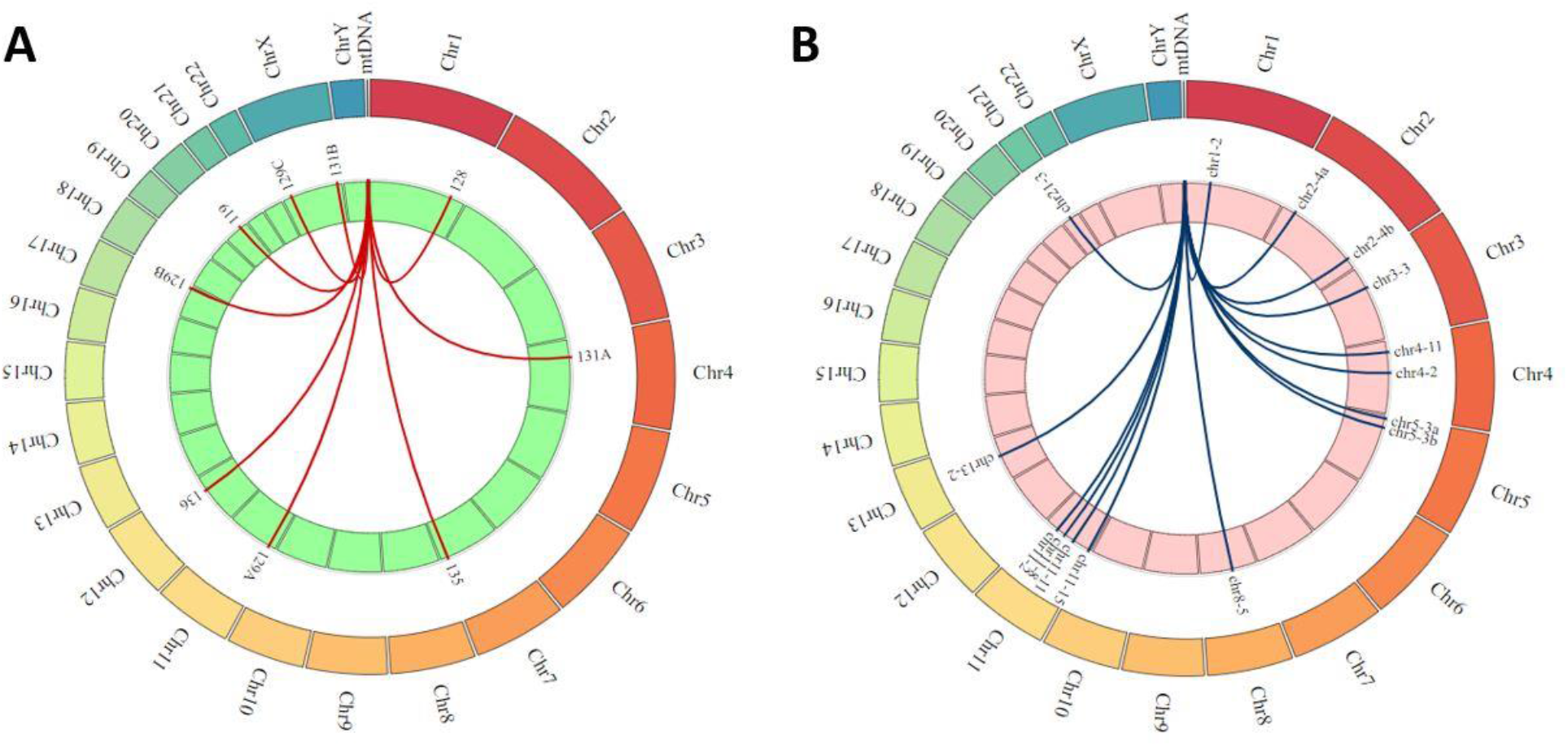
(A) Circos plot representing insertion location of S-*NUMT* s across chromosomes as identified by DINUMT. (B) Circos plot representing insertion location of G-*NUMT*s across chromosomes as identified by DINUMT.

**Table 1:** List of somatic mutations identified by DINUMT with their chromosomal location, length of the mitochondrial insert and patient no. \**NUMT*s were designated alphabetically when a single patient harbour more than one S-*NUMT*s.

| Chr. No. | Insertion Start | Insertion Stop | NUMT length | Patient No. |
| --- | --- | --- | --- | --- |
| 20 | 45709144 | 45709146 | 3895 | 119 |
| 1 | 210731831 | 210731888 | 6770 | 128 |
| 11 | 17491095 | 17491097 | 5087 | 129A* |
| 17 | 77335608 | 77335610 | 4304 | 129B* |
| X | 22480071 | 22480095 | 1933 | 129C* |
| 4 | 47023035 | 47023211 | 5436 | 131A* |
| X | 144335757 | 144335899 | 559 | 131B* |
| 7 | 140921797 | 140921880 | 5958 | 135 |
| 12 | 94944428 | 94944538 | 585 | 136 |

A total of 78 G-*NUMT*s were identified in 25 patients **(Figure 1B, Table 2)**. We named the G-*NUMT*s as follows: *chr-NO.-occurrence*. Five recurrently reported G-*NUMT*s present in chromosome 11 (2nos; chr-11-15, chr-11-11) chromosome 8 (chr-8-5), chromosome 4 (chr-4-11) and chromosome 2 (chr-2-4) were selected for confirmatory assay. Additionally, no previously reported *NUMT*s were found in the neighbouring genomic region (1kb upstream to 1kb downstream of the predicted insertion site) from each of the identified G-*NUMT* sites by UCSC genome browser with the *NUMT* track.

**Table 2:** List of germline mutations identified by DINUMT with their chromosomal location, length of the mitochondrial insert, frequency in the patients and the patients harbouring those *NUMT*s. ^#^ length of the longest *NUMT* observed in any of the patient carrying that *NUMT* is reported. ^*^length of the *NUMT* could not be determined by DINUMT.

| Chr. No. | Insertion Start | Insertion Stop | *NUMT* Length<sup>#</sup> | Frequency | Patient No. |  |  |  |  |  |  |  |  |  |  |  |  |  |  |
| --- | --- | --- | --- | --- | --- | --- | --- | --- | --- | --- | --- | --- | --- | --- | --- | --- | --- | --- | --- |
| 11 | 4656342 | 4656344 | 44 | 15 | 113 | 116 | 119 | 120 | 121 | 122 | 125 | 126 | 129 | 130 | 133 | 134 | 138 | 139 | 140 |
| 4 | 29441246 | 29441248 | 1353 | 11 | 116 | 117 | 119 | 120 | 121 | 122 | 125 | 126 | 137 | 139 | 140 |  |  |  |  |
| 11 | 49883568 | 49883572 | 377 | 11 | 113 | 117 | 119 | 122 | 130 | 133 | 134 | 135 | 137 | 138 | 140 |  |  |  |  |
| 11 | 77213043 | 77213050 | 39 | 8 | 113 | 120 | 121 | 122 | 125 | 126 | 138 | 139 |  |  |  |  |  |  |  |
| 8 | 63015422 | 63015431 | 121 | 5 | 116 | 119 | 122 | 129 | 133 |  |  |  |  |  |  |  |  |  |  |
| 2 | 33892478 | 33892482 | 130 | 4 | 121 | 134 | 137 | 140 |  |  |  |  |  |  |  |  |  |  |  |
| 2 | 214283781 | 214283783 | 37 | 4 | 119 | 134 | 139 | 140 |  |  |  |  |  |  |  |  |  |  |  |
| 3 | 56147220 | 56147222 | 63 | 3 | 117 | 119 | 139 |  |  |  |  |  |  |  |  |  |  |  |  |
| 5 | 8255946 | 8255948 | 43 | 3 | 116 | 119 | 138 |  |  |  |  |  |  |  |  |  |  |  |  |
| 5 | 32338582 | 32338584 | 2198 | 3 | 116 | 130 | 138 |  |  |  |  |  |  |  |  |  |  |  |  |
| 21 | 23080617 | 23080619 | * | 3 | 113 | 122 | 140 |  |  |  |  |  |  |  |  |  |  |  |  |
| 1 | 59661551 | 59661553 | 18 | 2 | 113 | 133 |  |  |  |  |  |  |  |  |  |  |  |  |  |
| 4 | 83616162 | 83616164 | * | 2 | 120 | 121 |  |  |  |  |  |  |  |  |  |  |  |  |  |
| 11 | 100015736 | 100015750 | 60 | 2 | 126 | 133 |  |  |  |  |  |  |  |  |  |  |  |  |  |
| 13 | 64162211 | 64162216 | * | 2 | 133 | 134 |  |  |  |  |  |  |  |  |  |  |  |  |  |

In 4 (119, 129C, 131B, 136) out of the 9 potential S-*NUMTs*, PCR-amplicons of length spanning the respective S-*NUMT* and the buffer region was observed from both tumour and matched blood. In two potential S-*NUMTs* (128, 131A), amplification of only buffer region was observed from both tumours and blood. No amplicons were observed for rest of the three potential S-*NUMTs* (129A, 129B, 135).

G-*NUMT* chr2-4 was detected in all 4 patients by PCR. G-*NUMT* chr8-5 was found to be present in blood and tumour DNA in only one patient. Presence of G-*NUMT* chr4-11 was detected in 3 OSCC-GB patients. Only three patients carried G-*NUMT* chr11-11. G-*NUMT* chr11-15 was detected in all 15 patients by PCR. Seven control individuals investigated from G-*NUMT* chr11-15 by PCR were positive for the specific amplicon. None of the G-*NUMT*s identified here interfered with ORF of any nuclear genes.

## Discussion

In 25 OSCC-GB patients investigated here, none harbour somatic *NUMT*s. Out of the 9 potential S-*NUMT* candidates identified by computational approach, none of them could be verified by PCR as somatic events. We have identified a number of germline *NUMT*s in oral cancer patients. But our study lacks the sample size required for estimating association of the observed G-*NUMT*s with oral cancer. The only G-*NUMT* recurrently observed in our sample set (chr-11-15) was also found to be present in blood DNA derived from non-cancer individual.

In a recent study, *Ju et al., 2015* have observed S-*NUMT*s in 12 breast cancer samples. Among these, only 10 are primary patient samples out of a total of 559 samples harbouring various types of tumours (ALL, AML, breast cancer, lung cancer, colorectal cancer, RCC, melanoma, glioma, prostate cancer etc.) studied in the TCGA project. These events could not be correlated with any clinical phenotype of cancer and were considered as passenger events. Our result, combined with the reports previously mentioned, indicates that S-*NUMT*s are ambiguous in nature and probably they occur in certain specific cancers (Ju et al., 2015; Yuan et al., 2017) and not others e.g. gingivobuccal oral squamous cell carcinoma.

To conclude, we did not find any OSCC-GB specific *NUMT*. Whether any or all of the G-*NUMT*s detected by u are novel polymorphic *NUMT*s remain to be verified in large data set. Our study of germline *NUMT*s in these patients did not reveal any evidence for association with OSCC-GB.

## Acknowledgements

We acknowledge the ICGC India Project team and the Core Technologies Research Initiative (CoTeRI) of NIBMG for generation of the sequencing data.

